# Natural variation in regulatory code revealed through Bayesian analysis of plant pan-genomes and pan-transcriptomes

**DOI:** 10.1101/2025.10.27.684862

**Authors:** Wei Wei, Xing Wu, Chandler A. Sutherland, Yuting Lin, China Lunde, Moises Exposito-Alonso, Ksenia Krasileva

## Abstract

Understanding the genetic code of cis-regulatory elements (CREs) is essential for engineering gene expression and modulating agronomic traits in crops. In plants, CREs underlying rapid evolution of gene expression often overlap with structural variation in promoters, making them undetectable using single-reference genomes. Here, we develop K-PROB (K-mer-based in silico PROmoter Bashing), a computational tool that learns from intraspecies promoter sequence and gene expression variation in pan-genomes and pan-transcriptomes to identify CREs controlling gene expression. K-PROB deploys a k-mer-based Bayesian variable selection framework to prioritize causal variable identification. We demonstrate the effectiveness of our approach in maize and soybean, two staple crops species. Applying K-PROB to genes with the most highly variable promoter sequences and the most diverse patterns of expression, such as nucleotide-binding leucine-rich repeat receptors, we identified k-mers enriched for bona fide transcription factor binding sequences, and overlapping with open chromatin regions and DAP-seq binding sites. Notably, multiple significant k-mers are located within presence/absence structural variants, highlighting structural variation in promoters as key drivers of transcriptional diversity of highly variable genes. We further validated the regulatory effects of identified k-mers on gene expression using luciferase reporter assays. Our results showcase a high-throughput and pangenomic approach for probing natural intraspecies cis-regulatory diversity, discovering new causative cis-elements, and facilitating future expression engineering across plant species.

**Significance Statement:** Understanding which DNA sequences control gene expression is essential for crop improvement. Current methods for identifying regulatory elements rely on expensive, specialized biochemical datasets typically limited to a single genotype. We developed a computational tool that links natural sequence variation and gene expression variation to identify functional regulatory sequences. Our tool employs a statistical framework that prioritizes causality over correlation, in contrast to most genome-wide association studies. Applying it to maize and soybean, two staple crops, we uncovered known and novel regulatory elements and validated them with molecular assays. Our approach is scalable, cost-effective, and efficiently utilizes natural variation from existing pangenomic datasets, opening new avenues for future crop engineering and studying gene regulation in diverse plant species.

## Introduction

Precise gene expression modulation has shown great promise in improving agricultural productivity and climate resilience in crop species (1, 2). It requires a mechanistic understanding of the spatiotemporal control of gene expression. The genetic information controlling expression is typically encoded in <u>c</u>is-regulatory <u>e</u>lements (CREs), such as transcription factor <u>b</u>inding <u>s</u>ites (TFBSs), and in <u>c</u>is-regulatory <u>m</u>odules (CRMs), which are assemblies of CREs that include core and proximal promoters, enhancers, silencers, and insulators (3). Identifying CREs or CRMs is a critical first step toward precise expression modulation through genome editing or synthetic biology approaches (1, 2). Traditionally, CREs within a promoter are identified through experimental promoter bashing, a process involving serial deletions of the promoter sequence followed by expression reporter assays (4, 5). While experimental promoter bashing can reveal CREs, it remains a low-throughput and labor-intensive approach, severely limiting the genome-wide discovery of CREs.

Over the past decade, a wide array of biochemical assays has been developed for high-throughput identification of CREs at the genome scale (3). <u>Ch</u>romatin immuno<u>p</u>recipitation-sequencing (ChIP-seq) and <u>D</u>NA <u>a</u>ffinity <u>p</u>urification sequencing (DAP-seq) profile transcription factor (TF)-DNA binding *in vivo* and *in vitro*, respectively, for TFBS discovery (6, 7). In addition to direct measurement of TF-DNA binding, TF-binding CREs or CRMs are usually located at nucleosome-depleted, accessible chromatin regions, so chromatin accessibility data from <u>a</u>ssay for transposase-<u>a</u>ccessible <u>c</u>hromatin with high-throughput sequencing (ATAC-seq) provides valuable information for CRE identification (8). When combined with single-cell technologies, ATAC-seq can further reveal cell-type-specific CRMs, enabling higher-resolution CRE identification at the spatial level (9). However, these biochemical assays typically require high sequencing depth and specialized protocols, resulting in such datasets being available only in a few plant model species and only in a single reference genotype, leaving the intraspecific cis-regulatory diversity unexplored.

Recent advances in bioinformatic and machine learning algorithms have offered multiple scalable approaches for CRE identification. Comparative genomics analyses have identified conserved noncoding sequences across different species, suggesting their evolutionary constraints and putative functional DNA sequences, such as CREs (1, 10). <u>TF b</u>inding sequence <u>m</u>otifs (TFBMs) can be used to predict putative CREs in other genotypes or species, due to the deep conservation of TF binding preferences among orthologues across plant lineages (11). These sequence-based methods tend to lead to false discoveries because the presence of CRE sequence does not equal regulatory function (8, 12). To address this, recent deep learning models such as DeepCRE and PhytoExpr build a sequence-to-expression relationship to annotate functional CREs. It attempts to learn general regulatory syntax by training on representative genomes and transcriptomes from multiple species (13, 14). These frameworks perform well for cross-species prediction, capturing conserved and variable regulatory motifs, but whether they can detect within-species cis-regulatory variation underlying expression variation is still uncertain.

Intraspecies cis-regulatory variation is a critical component for understanding diversity in gene expression and phenotype (15–17). In maize, a substantial proportion of quantitative trait loci (QTLs) for diverse traits lie within noncoding and open chromatin regions (17). Recent studies profiling TFBS variation across maize genotypes revealed that single-nucleotide polymorphisms (SNPs) and structural variants (SVs) at TFBSs explain much of the transcriptional diversity and heritable phenotypic variation in agronomic traits (15, 16). Orthologous genes within a species have been shown to differ in their degree of expression variability. A barley pan-transcriptome study showed that genes exhibiting more presence/absence variation, copy number variation, and other types of sequence-level diversity, which are often linked to rapidly evolving regions in the genome, tend to show highly variable gene expression too (18). For example, nucleotide-binding leucine-rich repeat receptors (NLRs) are rapidly evolving gene families involved in pathogen defense (19–21), and recently they were also shown to have extensive intraspecies allelic expression variation in addition to their well-known coding sequence diversity (21). Other examples of genes with intraspecies expression variation are also environment-responsive, such as the C-repeat/DRE-binding factor genes (*CBF*s) correlated with frost tolerance in barley (18) and the maize flowering time regulator APETALA2-like gene (*ZmRap2*.*7*) with the causal regulatory variation identified at the QTL *Vgt1* locus (16, 22). Despite these findings, systematic characterization of expression variation remains limited in most plant species, and the underlying cis-regulatory mechanisms, especially in highly variable genes, are still poorly understood due to gaps in available data and analytical approaches.

A common approach to study intraspecies regulatory variation is genome-wide association mapping for expression QTLs (eQTLs), which have been widely applied in diverse plant species (23–25). This method uses natural intraspecies variation from population-level genome re-sequencing data to identify variants significantly associated with gene expression. However, eQTL approaches have several limitations: they rely on short-read sequencing, read mapping to a selected reference and gene expression association with nearby tagging SNPs, which limits the identification of causal regulatory variants in genomic regions rich in structural variation (26, 27). Also, it does not reveal shared regulatory elements or regulation patterns across genes, as the analyses are conducted on a per-gene basis.

With advances in long-read sequencing technology, the rapid expansion of high-quality pan-genome and transcriptome data across both model and non-model plant species has facilitated the decoding of highly diverse genomic regions (28). Here we present K-PROB (K-mer-based in silico PROmoter Bashing), a computational tool that utilizes pan-genomic resources to identify causal cis-regulatory variants in highly variable genes. We deployed a k-mer-based approach to incorporate the intraspecies allelic diversity, CRE homology across genes, and a Bayesian variable selection model that prioritizes causality. Using maize and soybean data as proof-of-concept, K-PROB successfully identified known TF binding motifs without relying on prior TF-DNA binding information and pinpointed functional TF binding site variations embedded within large SVs. Contrasting motif enrichment or eQTL mapping approaches, the unique advantage of K-PROB to accurately identify the sequences of causal CREs, especially in promoters with pervasive SVs, makes it a valuable complement to existing tools for CRE discovery. Moreover, the highly variable genes on which K-PROB demonstrated strong statistical performance are enriched for stress responses, highlighting its potential to accelerate the engineering of key agronomic traits in crops.

## Results

### Pangenes with highly variable promoter and expression are enriched for environmental response

To systematically characterize intraspecies variation, we conducted a pangenomic analysis of gene expression and promoter sequence variation in maize and soybean. We used the maize pan-genome and RNA-sequencing (RNA-seq) data from eight tissues of 26 genetically diverse inbred lines (29) and a soybean pan-genome and RNA-seq data of leaf tissue for 27 lines (30, 31). We used published pangene membership to evaluate allelic variation where a pangene is defined as the collection of gene models sharing high sequence similarity and the same genomic locus in the pan-genome (32). We calculated the coefficient of variation (CV) of transcript abundance (defined as log_2_ transcripts per million, log_2_(TPM+1) for each pangene as a measure of the expression variation. The CV was further normalized to decorrelate it from mean expression as reported previously (Fig. S1, see Methods for details) (33). We used the 2kb sequences upstream of annotated genes as a proxy for promoter regions and calculated the promoter sequence similarity for each pangene by using a k-mer-based, alignment-free method, as it performs more accurately and efficiently for dissimilar sequences compared to sequence-alignment tools (34). The similarity score is defined as the average number of shared k-mers between any two promoters of a pangene.

After filtering out small-sized and silent pangenes (gene number <6 or mean log2(TPM+1) < 1), we kept 15,163 pangenes out of 78,456 for maize middle leaf tissue and 31,848 out of 64,835 for soybean leaf tissue. These pangenes exhibited a wide range of promoter sequence and expression variation. The log_2_ promoter similarity ranged from 4.9 to 10.9 with a median of 9.7 (IQR:9.2-10.2), and the normalized log_10_[CV^2^] ranged from -1.5 to 1.6 with a median of -0.0074 (IQR: -0.24-0.30) (Fig 1A). We investigated the functional annotations of pangenes that were in the top 30% for both expression and promoter sequence variation, as well as those in the bottom 30% for both features. GO analysis showed that the highly variable (hv) pangenes were significantly enriched for multiple biological processes related to environmental stress (One sided Fisher’s exact test, p-value < 0.05) including nutrients, water, and pathogens, such as “response to nitrate”, “root development”, “response to hydrogen peroxide”, “phenylpropanoid biosynthetic process” and “defense response” (Fig 1B, Dataset S1). In contrast, the least variable pangenes were enriched for primary cellular processes such as respiration, gene transcription, protein metabolism, and methylation (Fig. 1B, Dataset S1). Pfam domain enrichment analysis of the hv pangenes revealed many domains found in proteins mediating cellular stress responses, including leucine-rich repeat and NB-ARC domains typically seen in immune receptors, Glutathione S-transferases (GSTs) that scavenge reactive oxygen species, and dirigent-like proteins that modulate cell wall for stress and root nutrient transport (one-sided Fisher’s exact test, p-value < 0.05); Fig. 1C) (20, 35–37). Soybean pangenes also exhibited a wide range of intraspecies variation. The log_2_ promoter similarity ranged from 6.3 to 10.9 with a median of 10.6 (IQR:10.5-10.8) and the normalized log_10_[CV^2^] ranged from -1.4 to 1.9 with a median as 0 (IQR: -0.26-0.30) (Fig S2). Similar GO and Pfam domain enrichment related to environmental response were observed in soybean hv pangenes (Fig S2), suggesting a consistent pattern of intraspecies expression variation across plant species.

**Figure 1.**
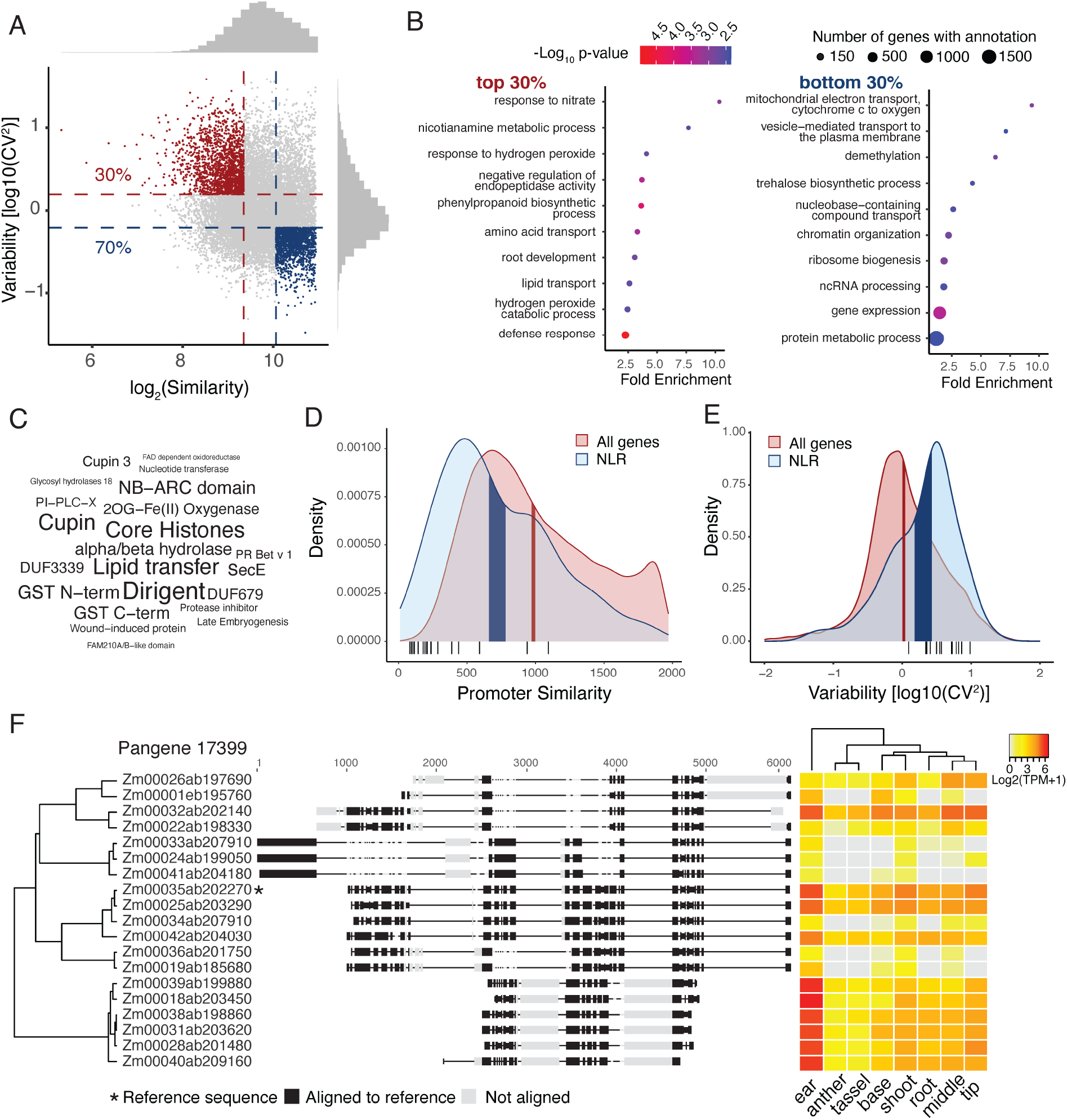
Pangene variation analysis and functional category enrichment. (A) Distributions of pangene promoter similarity and expression variation. The promoter similarity indicated by log_2_(similarity) and expression variation indicated by log_10_[coefficient of variation (CV)^2^] measure the allelic variation among genes within pangene. Pangenes in the 30% most variable and 30% least variable for both promoter and expression distributions were marked by red and blue. Histograms show distributions of individual variables. (B) GO term enrichment for top and bottom 30% pangenes. Displayed GO terms are significantly enriched at a cutoff of p-value < 0.05 for one-sided Fisher’s exact test. (C) Pfam domain enrichment for top 30% pangenes. Only domains assigned to 3 or more pangenes in the top 30% and with enrichment p-values < 0.05 for one-sided Fisher’s exact test were reported. The size of the text is proportional to the number of observations of that domain in the top 30%. (D-E) The distributions of NLR pangenes and all pangenes for promoter similarity (D) and expression variability (E). Shaded areas denote the 95% confidence interval of the bootstrap mean (n=10,000). Black bars underneath the density plot represent the subset of NLR pangenes reported to have extremely hv protein sequences. (F) The 2kb promoter sequence alignment and expression in multiple tissues of an example NLR pangene. Hierarchical clustering on the left is based on the pairwise similarity scores output from the SV-aware sequence alignment tool Mauve, with the starred promoter as the alignment reference.

The plant nucleotide-binding leucine-rich repeat receptor (NLR) family contains immune receptors that typically exhibit abundant intraspecies protein level diversity, including subsets of genes that are rapidly evolving to enable new pathogen recognition (21). We found that at the expression level, NLR pangenes also displayed significantly higher promoter sequence and expression variation compared to all other pangenes in the maize genome (Fig 1D,E; bootstrap mean test, n=10,000). This higher variation was also observed for NLR expression in other maize tissues (Fig S3A) and in soybean (Fig S3B). A subset of NLR pangenes with highly variable protein sequences also tend to show hv promoter sequences and expression in general (Fig 1D,E) (21). As an example of a hv NLR pangene, the pangene pan17399 showed extensive variation in promoter sequence alignment, which is exemplified by indels and structural variants (SVs) ranging from a few base pairs to nearly a kilobase (Fig 1F). Moreover, we clustered promoters for this pangene by sequence similarity and observed a general correlated pattern between promoter variation and expression based on visual inspection, suggesting these sequence variations might be the cause of expression changes (Fig 1F).

### Bayesian joint modeling of intraspecies cis-regulatory variation using k-mers

Given that highly variable (hv) pangenes exhibited correlated patterns of promoter sequence and gene expression variation (Fig 1F), we hypothesize that association analysis can be performed to identify functional sequence variants. However, due to pervasive SVs in these promoters, pinpointing the causal cis-regulatory elements (CREs) would be challenging using traditional association methods that rely on sequence alignment to a single reference genome and use SNPs as markers, such as most eQTL mapping studies (23–27). To facilitate systematic identification of CREs in pangenes with highly diverse promoter sequences, we developed K-PROB (K-mer based in silico PROmoter Bashing), which uses k-mers extracted from promoter regions in fully-assembled pan-genomes and associates k-mer counts with gene expressions from pan-transcriptomes, removing reliance on any single reference genome (Fig 2A). Also, unlike SNPs that represent unique genomic positions, k-mers can be shared across different pangenes and can have lengths matching typical transcription factor binding motifs (6-12 bp) (3). This allows common regulatory patterns across pangenes to be learned and enables identified k-mers to directly represent TF binding sequences. To increase the detection power and biological relevance, we also grouped similar k-mers into clusters as most TF binding allows mismatches (11). For the statistical model, K-PROB jointly models all k-mer clusters and expression of all pangenes using a Bayesian variable selection regression method. This joint model could yield more accurate estimation of kmer clusters with non-zero effects and thereby prioritizes causative CRE identification, in contrast to single-variable marginal regression methods commonly used in GWAS models (38). The <u>p</u>osterior inclusion <u>p</u>robability (PIP) of each k-mer cluster was estimated and those with high PIP were considered as putative CREs (details in Material & Methods). However, occurrences of CRE sequences do not necessarily equate to functional TF binding sites (8, 12). To identify functional sites for each high-PIP k-mer cluster, K-PROB then calculates the effect of each k-mer cluster on a per-pangene basis, and only keeps k-mer cluster - pangene pairs with significant effects. The kept k-mer cluster sites are considered functional (Fig 2A, the last panel). In summary, we expect that the intraspecies variation between promoter alleles, CRE homology across pangenes, and the Bayesian variable selection framework of statistical modeling all increase the likelihood of causal CRE identification, and distinguish our method from other existing tools.

**Figure 2.**
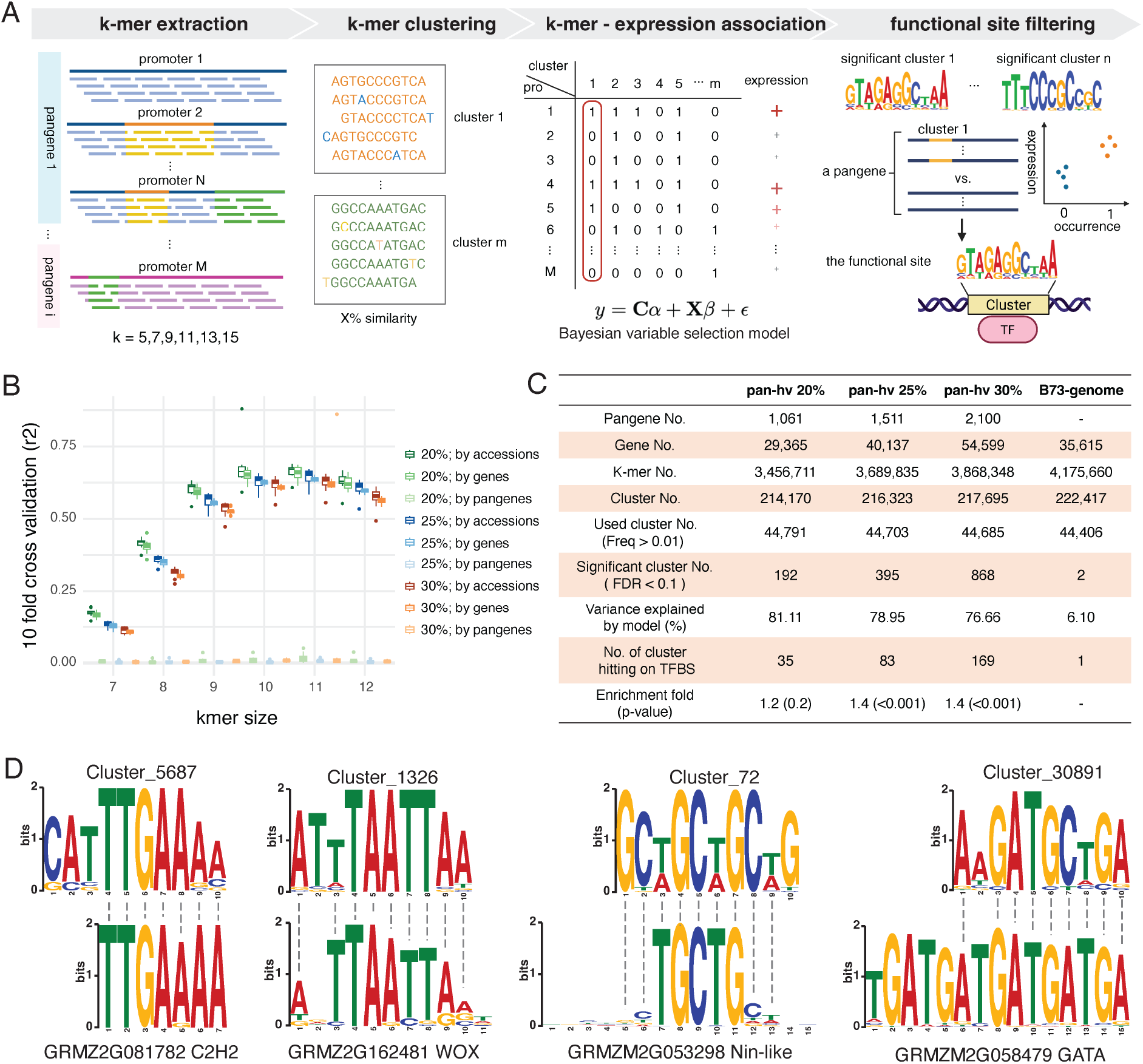
K-mer based *In silico* promoter bashing identified k-mer enriched for known TF binding motifs. (A) Schematic representation of the K-PROB framework, which uses promoters and expressions from pan-genomes and pan-transcriptomes as the sole input to identify CREs and the corresponding intraspecies CRE variations responsible for expression variations. (B) Evaluation of the model performance across different k-mer sizes by 10-fold cross-validation. The legend on the right demonstrates the three datasets of pangenes in the top 20%, 25%, and 30% for both promoter and expression variation, and different strategies to divide training and testing sets for cross-validation. (C) Summary metrics from running K-PROB on different maize datasets and enrichment test of significant k-mers for known TF binding motifs. The four datasets include the top 20%, 25%, and 30% most variable pangenes and all genes from the single B73 genome. (D) Sequence similarity between motifs computed from significant k-mer clusters and known TF binding motifs from the PlantTFDB. Shown motifs are the most frequent ones among the top 30% highly variable pangenes.

As k-mer size is a critical parameter that could affect the model fitting and statistical power for candidate identification, we first decided which k-mer size to use. To evaluate the response of different k-mer sizes, we generated k-mer clusters with different k-mer sizes ranging from 7 bp to 12 bp from three datasets with various percentile cutoffs: pan-genes showing 20%, 25%, and 30% top variation for both promoter sequences and gene expression. Across all datasets, the total number of unique k-mers and k-mer clusters increased with k-mer size, whereas the percentage of k-mer clusters included in the model decreased (Table S1) because more low-frequency k-mers are filtered out by the cutoff requiring k-mer cluster to occur in more than 1% of total hv genes. With different k-mer sizes, we evaluated their gene expression prediction accuracies of our models using 10-fold cross-validation to assess the model fitting. We divided the training and testing dataset in three different ways: (1) by genes (2) by inbred lines (3) by pangenes. The former two dividing strategies showed a similar trend of predicted accuracies of k-mer size response: the accuracies were lowest at k = 7 where the R^2^ ≈ 0.12, increases as k-mer size increases and peaked at k = 11 where the R^2^ ≈ 0.62 (Fig 2B). We tested all three different datasets which all showed k=11 as the peak of the accuracy curve so we decided to use k = 11 for the rest of our analysis (Fig 2B). However, splitting data by pangenes for training and testing yielded low prediction accuracies for all k-mer sizes, suggesting this model can be used for predicting allelic promoters but is limited at predicting distinct promoters of unseen hv pangenes. Interestingly, we did the same k-mer size test using soybean pan-genome data and also obtained k=11 as the optimal k-mer size (Fig S4).

### K-PROB identifies significant k-mer clusters enriched for transcription factor binding motifs

We hypothesized that different subsets of highly variable (hv) pangenes may influence CRE identification, so we ran K-PROB using the maize pan-genome datasets with three variation thresholds: pangenes in the top 20%, 25%, and 30% for both promoter and expression variation. These datasets contained 1,061, 1,511, and 2,100 pangenes, corresponding to 29,365, 40,137 and 54,599 genes from 26 maize inbred lines, respectively (Fig 2C). Although the number of pangenes and genes varied substantially (up to a two-fold difference), the total number of unique k-mers (3.5– 3.8 million) extracted from promoters, total k-mer clusters (0.21–0.22 million), and occurrence-filtered clusters used for modeling (∼45,000) remained similar (Fig 2C). With comparable numbers of input k-mer clusters, K-PROB identified 192, 396, and 868 significant 11-mer clusters (FDR <= 0.1 based on PIP) for datasets with increasing pangene numbers, respectively. To show that intraspecies allelic variation is critical for effective identification of significant k-mer clusters, we also ran K-PROB using all the genes from one single B73 genome. Despite a similar number of input genes (35,615) and k-mer clusters (∼44,500), this run yielded only 2 significant clusters and the resulting model only explained ∼6% of the expression variation, compared to over 75% for all three datasets using 26 genomes and top hv pangenes (Fig. 2C).

To demonstrate that K-PROB can identify functional CREs and not just associated DNA sequences, k-mers in the significant clusters were aligned to known maize TFBMs from the PlantTFDB databases. The significant clusters obtained by 25% and 30% datasets are significantly enriched for k-mers with high sequence similarity to known TFBMs (1.4 fold, permutation test, n=1000, p<0.001) (Fig 2C), suggesting K-PROB is an effective tool to identify CREs when only being given promoter sequences and expressions datasets. We also ran K-PROB using 20%, 25% and 30% of highly variable pangenes in the soybean pan-genome and pan-transcriptome data, which also yielded k-mer clusters significantly enriched for TF binding sequences (Table S2). These results supported the generalizability of K-PROB application to different plant species. For the rest of the analyses, we continued with the maize 30% datasets as it identified the highest number of significant k-mer clusters with significant enrichment for TFBMs.

We investigated which TFs were represented by our identified significant k-mer clusters and thus could be involved in regulating maize hv pangenes. We computed motifs from significant k-mer clusters and aligned these motifs to putative maize TFBMs. As many clusters aligned to the same TFBMs, after removing duplicates, we kept 67 TFBMs from the PlantTFDB belonging to 29 TF families (Dataset S3). The three most frequent TFs are a C2H2 TF zinc finger protein 10 (GRMZM2G081782), a Nin-like TF (GRMZM2G053298) and a WUSCHEL homeobox 19 (WOX19) TF (GRMZM2G162481), and they were aligned by significant k-mer clusters with total occurrences of 36,076, 19,517 and 17,818 times in the hv pangenes (Dataset S3 and Fig 2D). The Nin-like TF GRMZM2G053298 was shown to have expressions correlated with leaf size-related traits and its homologs in Arabidopsis functions in leaf development (39, 40). The WOX19 TF belongs to a plant-specific family WOX known to regulate genes critical for growth and development (41). Besides WOX19, the binding motif of WOX13a (GRMZM2G069274) was also aligned with two significant k-mer clusters corresponding to 4,666 total occurrences (Dataset S3). This TF was reported to modulate plant height and also respond to salt stress (41, 42). As one of the most frequent TFs in the list, ZML2 (GRMZM2G058479), a GATA family TF, had high binding motif similarity to four clusters all with negative effects on expression (Fig 2D, Dataset S2 and S3). ZML2 acts as a transcriptional repressor that inhibits lignin biosynthetic genes in maize and will be degraded by wound stress (43).

K-mer clusters that did not align to any known TFBMs may represent novel motifs absent from current databases. PlantTFDB predicts putative TFBMs for over 150 plant species by homology to Arabidopsis motifs (44), a strategy that can miss species-specific TFBMs. To test whether K-PROB recovers maize-specific TFBMs lacking in PlantTFDB, we cross-referenced our significant k-mer clusters with a recent DAP-seq-derived motifs for 200 maize TFs (16) and found 46 additional k-mer clusters with homology to maize TFBMs corresponding to 38 distinct TFs (Dataset S3), in addition to the 169 predicted by PlantTFDB. For example, cluster_289 aligned with ARF39, cluster_26910 with ABI8, and cluster_56596 with MYB131 (Fig. S5). These findings support K-PROB’s k-mer-based strategy for uncovering novel TFBMs, especially in non-model species where validated motif catalogs remain largely incomplete.

### K-PROB further filtered k-mer occurrences for functional regulatory sites

The presence of a TFBM sequence in the genome does not necessarily indicate true TF binding with regulatory activity, as it also depends on flanking sequences or chromatin context (8, 12). The effects of significant k-mer clusters (FDR <=0.1) estimated by the Bayesian variable selection model represent average effects across all their genomic occurrences, including both functional and non-functional. These estimated effects (defined as the change in log_2_ expression per occurrence of the sequence) were generally small, ranging from 0.056 to 0.31 for potential activators and from -0.079 to -0.24 for potential repressors (Dataset S2; Fig 3A, the heatmap on the left). To further filter for functional sites for each k-mer cluster, we calculated its effect on a per-pangene basis, by performing a linear regression between its occurrence and gene expression within each pangene. Only pangenes showing a significantly non-zero effect were kept as potential functional sites for the k-mer cluster (Fig. 3B). We hypothesize that k-mer sites retained through this functional filtering would represent true regulatory sites with greater confidence.

**Figure 3.**
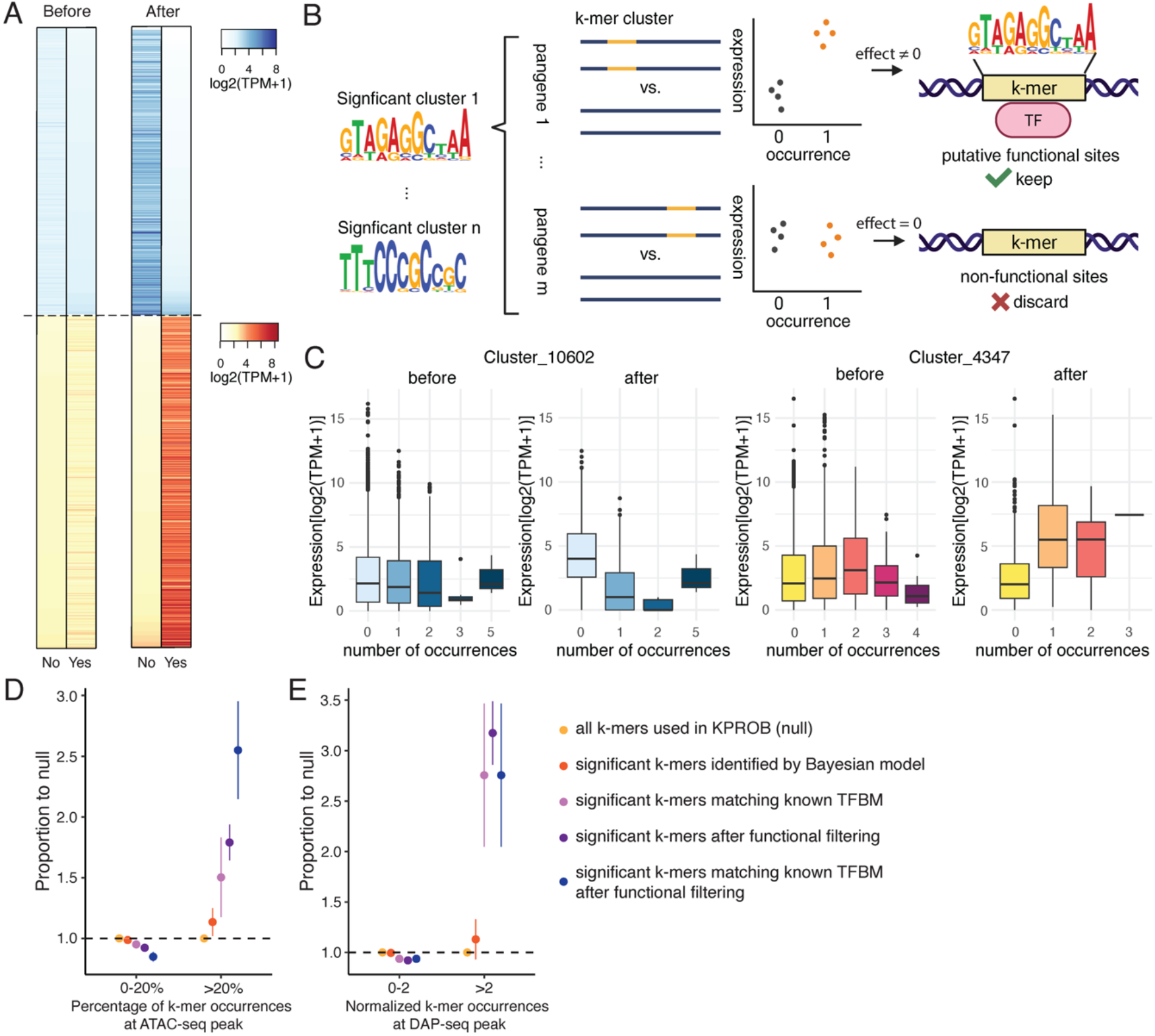
Evaluation of the impact of functional site filtering. (A) Average gene expression with and without the k-mer cluster before and after functional site filtering. “Yes” indicates presence and “No” indicates absence of the k-mer cluster. Each row of the heatmap represents a k-mer cluster and the color intensity indicates the average log_2_(TPM+1) of genes in the corresponding group. Only pangenes containing the k-mer cluster are included for each k-mer cluster. K-mer clusters with predicted negative effects are colored blue and positive effects colored red. Four heatmaps were sorted independently by the groups with expected lower average expression. (B) Overview of functional site filtering process. (C) Boxplots showing the effect of k-mer occurrences before and after functional site filtering for two representative k-mer clusters with predicted negative effect (left) and positive effect (right), respectively. Number of occurrences indicates the number of occurrences within each promoter. (D-E) Measurement of functional regulatory site enrichment at different stages of the K-PROB pipeline by comparing to ATAC-seq (D) and DAP-seq data (E), respectively. For each k-mer cluster, the percentage of k-mer occurrences at ATAC-seq peaks is calculated as k-mer occurrences under narrow peaks divided by total occurrences in all promoters. Normalized k-mer occurrences at DAP-seq peak is calculated by aggregated k-mer occurrences under DAP-seq narrow peaks of all TFs, divided by total occurrences in all promoters. The error bar represents the standard error.

K-PROB identified significant k-mer clusters across a broad set of pangenes, with each cluster detected in 47 to 1,039 pangenes, and a median of 135 (Fig. S6A). After functional filtering, 2% to 25% of the pangenes were kept for each k-mer cluster, with a median of 13% (Fig. S6B and Dataset S4). By comparing the average k-mer cluster effects before and after filtering, we observed a general increase in effect size in both positive and negative directions (Fig. 3A). This was reflected by stronger expression differences between “presence” and “absence” of the k-mer sequence after filtering (Fig. 3A). For example, the estimated effect of Cluster_4347, representing a putative activator binding sequence, increased from 0.12 to approximately 2 (Fig. 3C). Similarly, the effect of Cluster_10602, representing a putative repressor binding sequence, had the magnitude increased from -0.14 to approximately -2 (Fig. 3C).

We next evaluated whether our functional site filtering approach successfully identified functional regulatory sites by comparing the results to other independent functional annotations. As functional CREs are often located in open chromatin regions, we first investigated how many of our significant k-mer cluster occurrences overlapped with open chromatin regions, using the ATAC-seq data of the 26 inbred lines (29). For each k-mer cluster, we calculated the percentage of its occurrences falling under ATAC-seq peaks and then binned the percentages into 0-20% and >20% groups, where the latter indicates the high-confidence group for functional sites. Compared to the null distribution of all k-mer clusters included in the Bayesian variable selection model, the identified significant k-mer clusters showed a modest enrichment in the high-confidence group (1.14-fold, 95% CI: 0.91-1.37) (Fig. 3D). More importantly, conducting functional site filtering with these significant k-mer clusters further increased the enrichment to 1.79-fold (95% CI: 1.52-2.07), supporting the effectiveness of our functional site filtering strategy. Performing functional filtering on significant k-mer clusters matching known TFBMs yielded the highest enrichment in the high-confidence group (2.57-fold 95% CI: 1.81-3.38). To provide another orthogonal source of evidence, we also counted the overlaps between our identified k-mer cluster sites and DAP-seq peaks from 200 maize TFs (16). Notably, applying functional site filtering to significant k-mer clusters identified by the Bayesian variable selection model yielded the highest enrichment (3.19-fold, 95% CI: 2.55-3.84), even more than combining the information from known TF binding motifs (2.7-fold, 95% CI: 1.43-4.3, Fig 3E), further supporting our functional filtering approach.

### Newly identified causal CRE variations are supported by experimental validation

By performing association with the Bayesian variable selection model and subsequent functional site filtering, we found that K-PROB can identify regulatory sites containing natural intraspecific variation that were not previously reported from functional assays. For instance, K-PROB identified two k-mers from Cluster_72 that match a Nin-like family transcription factor (TF). These k-mers potentially mediate gene expression variation of the pangene pan09953, which encodes a calcium-dependent lipid-binding (CLB domain) family protein. These two k-mers are located in a highly polymorphic promoter region in an open chromatin state, and the presence of either could significantly boost the gene expression (Fig. 4A). Although little is known about maize CLBs, in Arabidopsis, CLB (AT3G61050) has been reported to act as a repressor of stress response (45). The observation that a growth-regulating TF mediates the expression of CLB in maize suggests a potential transcriptional regulatory node balancing plant growth and stress response. The identified CRE variation could be responsible for natural phenotypic variation underlying the growth-resilience trade-off in plants (46). Additionally, a k-mer of Cluster_4347, which shows homology to the WOX13a binding motif, is located within a ∼300bp insertion in several promoter alleles of pan08252, overlapping with a ∼750bp ATAC-seq peak. Pan08252 encodes an uncharacterized membrane protein containing a lipoprotein lipid attachment site domain (Fig 4A). Similarly, a k-mer of Cluster_11172 that matches a heat shock factor binding motif was detected within a ∼50bp insertion in promoters regulating pan02430 that encodes a bHLH148 TF (Fig. S7). These examples further support the role of structural variation as a major driver of transcriptional diversity of these hv pangenes. In addition, although these k-mers have sequence homology to known TF binding motifs, K-PROB uncovered regulatory relationships between the TFs and the regulated hv variable genes and revealed previously unknown CRE variants that contribute to the intraspecies expression variation.

**Figure 4.**
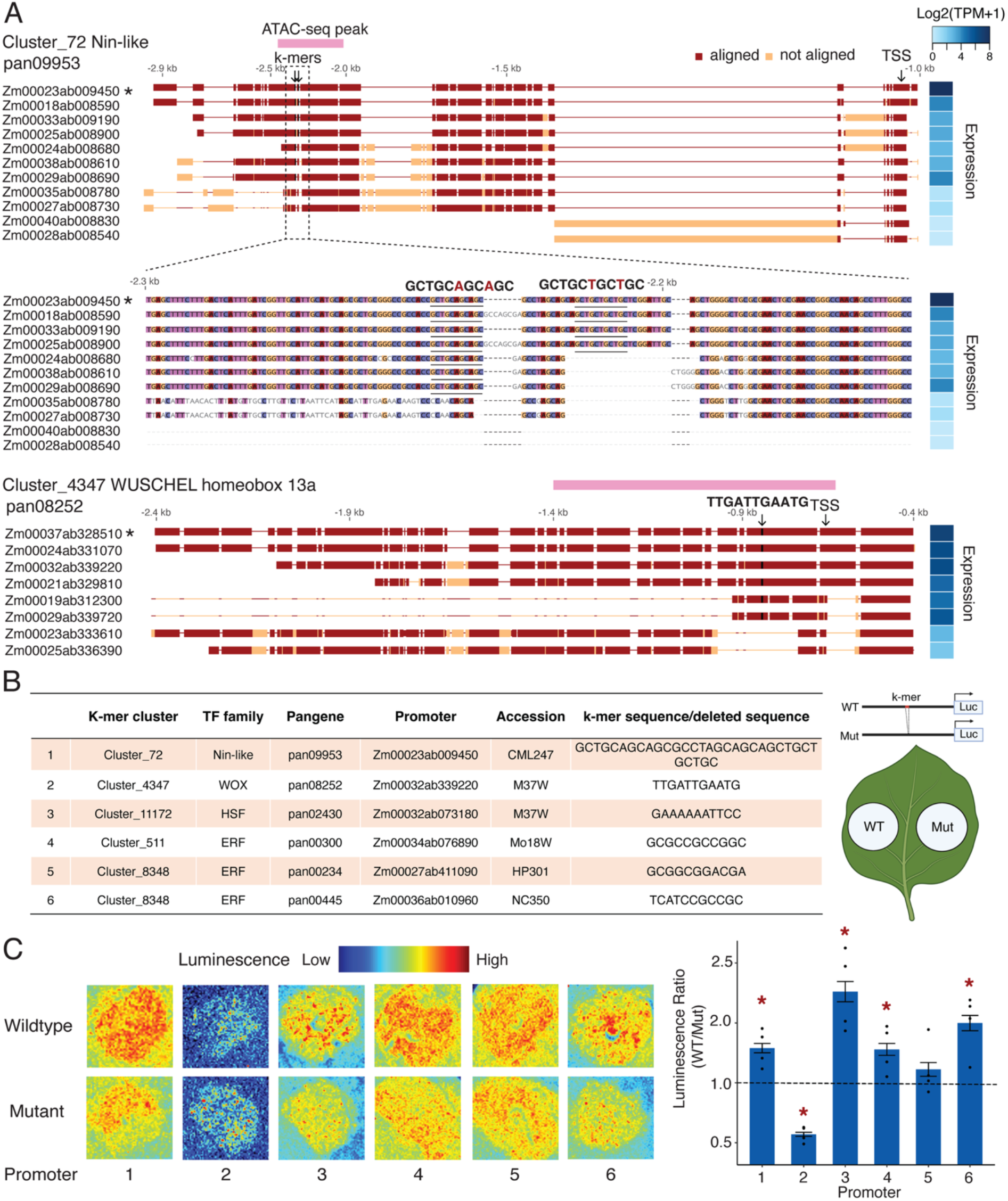
K-PROB pinpointed previously unknown cis-regulatory element (CRE) variants responsible for expression variation. (A) Examples of three significant k-mer clusters located within highly polymorphic promoter regions or structural variants (SVs). Promoter sequence alignments were performed using the SV-aware alignment tool Mauve, with the starred promoter used as the alignment reference. TSS: predicted transcription start site from Softberry. ATAC-seq peak information was obtained from Hufford et al. (2021) (29). Letters above the alignments indicate the identified k-mer position and sequence. (B) k-mer clusters selected for functional validation using a promoter-luciferase reporter assay in *Nicotiana benthamiana* (on the left) and the leaf infiltration layout (on the right). WT refers to native maize promoters containing the k-mer sequence, while mutant indicates promoters where only the k-mer sequence was removed. (C) Representative luminescence images and quantitative measurements from the promoter-luciferase reporter assay. Images and measurements were taken two days post *Agrobacterium* infiltration. To minimize variation in expression efficiency across leaves, luminescence ratios were calculated by dividing the luminescence of the WT construct by that of the mutant construct infiltrated on the same leaf. Red stars indicate ratios significantly different from one (dashed line), denoting a significant difference from WT (t-test, p-value < 0.05).

To experimentally validate the effects of these CRE variants, we conducted promoter luciferase reporter assays in a heterologous system of *Nicotiana benthamiana* using native maize promoters containing the k-mer sequence and mutant promoters where only the k-mer sequence were removed. We validated five k-mer clusters with estimated positive effects in expression, covering six potential cis-regulatory sites (Fig. 4B). Five out of six showed significant expression differences between the wild type and the mutant constructs, with fold ranging from 1.5 to 2.3 (Fig. 4C). Four k-mers exhibited effects in *N. benthamiana* that were consistent in direction with the estimated effects in maize. However, the k-mer from Cluster_4347 showed a highly significant but opposite effect in *N. benthamiana*. While this exception still supports the regulatory function of this site, it also suggests potential functional divergence of TFs with conserved binding profiles across species (47). In summary, our experimental validation supports that K-PROB successfully pinpointed causal CREs within large SVs.

## Discussion

Our study developed a novel framework, K-PROB, to identify causal cis-regulatory elements (CREs) using natural intraspecific promoter and expression variation using pan-genome and pan-transcriptome datasets. Pangenome-wide variation analysis revealed that pangenes with highly variable (hv) promoter sequence and gene expression levels are enriched for functions related to environmental responses in both maize and soybean, including stress tolerance, pathogen defense, and nutrient utilization. Our observation raises the question about the functional implications of the high allelic expression diversity, driven by elevated levels of cis-regulatory variation. Variation in expression can precede and promote protein coding sequence diversification, as previously observed in NB-ARC domain-containing NLR immune receptors, which were highly enriched in the top variable genes in our study (20). We also found significant enrichment of redox-related genes such as GSTs and thioredoxins, which play key roles in both stress response and development (48, 49). Although these genes are typically stress-inducible, the basal expression differences observed in steady-state leaves may indicate variation across genotypes in the priming capacity for stress responses, potentially contributing to differences in their ability to adapt to diverse growing conditions (18, 50). Additionally, expression variation in leaf tissue for nutrient-response genes, including those involved in nitrate response and assimilation, may underlie differences in leaf growth and metabolic rates across environments with varying nutrient availability, as nitrogen assimilation is essential for synthesizing enzymes and chlorophyll required for photosynthesis and other metabolic processes in leaves (51).

Overall, k-mer-based approaches have gained increasing attention in the pangenomics era due to their versatility in handling extensive sequence diversity across genomes without reliance on aligning sequences to a reference (34, 52). However, application of k-mer approaches to understanding variability in regulation of gene expression remain limited. In our study, we leveraged k-mer-based methods at multiple stages of our workflow as they present unique advantages over alignment-based methods. To compare hv promoter sequences, we used the alignment-free tool PanKmer, which represents pangenomic sequences based on the presence or absence of unique k-mers (34). In addition to being better suited for highly divergent sequences, k-mer-based comparisons are also significantly more computationally efficient than traditional multiple sequence alignment tools (34), enabling scalable analysis across tens of thousands of pangenes in our study. For K-PROB modeling, we used k-mer clusters as variables in a Bayesian variable selection model and showed that K-PROB can directly identify significant k-mer clusters as CREs, as evidenced by enrichment for k-mer sequences homologous to known CREs. By using k-mers in the size range of TFBSs, followed by clustering to capture mismatches, we better approximated natural TF binding sequences and improved causal identification. This illustrates a unique advantage of K-PROB over SNP-based association tools, which often identify correlational but not causal variants, and SV-based association tools, which may identify causal variation but lack the resolution to pinpoint the specific CREs within structural variants (26, 27). While k-mer approaches have been deployed in various forms of GWAS to associate natural genetic variation with traits, those studies typically use longer k-mers as unique genetic markers (53, 54). On the other hand, k-mers have also been applied in various CRE identification tools where they represent putative CREs, such as in machine learning classifiers trained on TF binding and non-binding sequences in human (55, 56), or in motif enrichment analyses based on co-expressed genes in Arabidopsis (57). K-PROB explores a new scenario that unifies the use of k-mers to represent both regulatory sequences and natural allelic variations in the promoter on a pan-genome scale, enabling the identification of causal CREs for highly variable genes in pan-genomes.

Although K-PROB proved to be an effective tool, several aspects can be improved or expanded in future research. First, this study used genomes from only ∼25 maize or soybean accessions, but as high-quality pangenomic datasets continue to grow (32), incorporating more intraspecific genetic and transcriptional variation is expected to greatly enhance statistical power and resolution. Additionally, the Bayesian variable selection model implemented in K-PROB is highly adaptive and can incorporate orthogonal biologically grounded functional annotations as prior information to further prioritize causal kmer cluster identification (58). The recent expansion of functional annotation resources such as DAP-seq and ATAC-seq (8, 16) might further boost its ability to pinpoint causal CRE variants. Second, our study, as well as most existing sequence-to-expression deep learning models for functional cis-variant prediction, focuses on steady-state gene expression in plants (13, 14). However, while genes involved in plant plasticity under diverse stress conditions have been widely identified (57, 59), population-level variation in stress-inducible expression remains poorly characterized (60). K-PROB offers a promising framework to link natural genetic variation with inducible expression variation for the identification of causal stress-responsive CRE variants. Third, K-PROB selects an optimal and a fixed k-mer length between 7-12bp based on model fitting specific to each datasets. While effective for current datasets, more flexible methods that accommodate variable k-mer lengths could further improve the power to identify TF binding motifs of diverse sizes. For example, a machine learning based CRE classifier gkm-SVM uses gapped k-mer features, which include non-informative gaps to allow longer k-mers (>15bp) (56). Finally, the current model only considered the additive effects of each k-mer (putative CRE) on gene expression whereas growing evidence across species, including plants, showed that co-localized TFs often act cooperatively in cis-regulatory modules to influence gene expression (16, 61). Future extensions of K-PROB could include interaction terms to capture these biologically meaningful TF-TF interactions and enhance its biological interpretability.

In summary, we present a new framework of CRE identification based on linking pan-genome with pan-expression variation using a k-mer-based approach and a Bayesian variable selection model that promotes causality. We further refine candidate CREs with functional site filtering and *in planta* validation to validate causal relationship. While this work can be expanded across new future datasets, its application to existing plant pan-genomics and pan-transcriptomic resources provides plant biologists and breeders alike with rational approaches for precise gene expression modulation.

## Materials and Methods

### Pan-genome and pan-transcriptome data collection and processing

The Maize pan-genome sequence and gene annotation data of 26 maize inbred lines were downloaded from MaizeGDB (https://www.maizegdb.org/NAM_project). The corresponding processed RNA-seq data (TPM) from different tissues was obtained from Prigozhin et al. 2024 (21) with the raw data generated by Hufford et al. 2021 (29). Soybean pan-genome sequence and gene annotation data of 27 soybean and *Glycine soja* accessions were downloaded from Genome Warehouse database in the BIG Data Center (https://bigd.big.ac.cn/gsa/index.jsp) under Accession Number PRJCA002030 (30). RNA-seq generated from fully expanded young leaves from 2-week-old seedlings was obtained from the Genome Sequence Archive in the BIG Data Center (PRJCA00936) (31). To process soybean RNA-seq data, quality trimming was performed using Trimmomatic v0.39, the quality of trimmed reads was checked with fastQC v0.12.1, reads were mapped using STAR v2.7.11 and the number of reads aligned were counted and normalized to TPM by RSEM v1.3.3 (62–64).

### Promoter sequence similarly and gene expression variation calculation

BED files recording genomic positions of 2kb upstream of annotated genes were created with a custom python script and sequences were extracted using bedtools v2.31.0 (65). Homolog pangene clustering information was obtained from MaizeGDB for maize and SoyBase for soybean (https://www.soybase.org/), both of which used Pandagma for clustering (66). The promoter sequence similarity of a pangene was computed by averaging the pairwise promoter similarity of all possible pairwise comparisons, which were calculated using PanKmer, a k-mer-based alignment-free sequence comparison tool as it has unique advantages for more dissimilar sequence alignment (34). The similarity score is defined as the number of unique 31-mers shared between two 2kb promoters so it ranges from 0 to 1970 and log_2_ similarity from 0 to 10.94. To measure gene expression variation of a pangene, log_10_ squared coefficient of variation (log_10_CV^2^) was calculated using expression values log_2_(TPM+1) of all genes. Because CV^2^ decreases with mean gene expression increases, as previously reported in humans (33, 67) and also observed in maize (Fig S1), to account for this bias and perform a fair CV comparison across pangenes with different expression levels, log_10_CV^2^ was further normalized by subtracting the expected log_10_CV^2^ for each pangene according to its mean expression level using a running median approach (33, 67).

### GO term and PFAM domain enrichment analyses

To identify highly variable pangenes for enrichment analyses, we only kept pangenes that are considered expressed (mean log2(TPM+1) > = 1) and pangenes having more than 5 gene members because we found those small-sized pangenes are enriched with very similar promoters, which could be a sign of inaccurate pangene clustering. The PANNZER2 annotations for the 26 NAM lines were downloaded from Maize GDB (https://download.maizegdb.org/GeneFunction_and_Expression/Pannzer_GO_Terms/) and the gff3 annotation files for the 27 soybean lines containing PANNZER2 and Pfam annotations were downloaded from soybase (https://www.soybase.org/). All GO terms assigned to genes within a pangene were then transferred to their pangene. GO term enrichment analysis was then performed using the R package TopGO (Alexa and Rahnenfuhrer 2023) version 2.56.0 and enrichment was calculated using the Fisher’s exact test and the “elim” algorithm. Only GO terms that were assigned to 3 or more pangenes of the test set and with enrichment P-values less than 0.05 were reported. Pfam annotations were downloaded from maize GDB (https://download.maizegdb.org/) and extracted from the soybean gff3 annotation files. Annotations were assigned to a pangene if at least half of the pangene members had that domain annotation. A one-sided Fisher’s exact test for enrichment was performed using the R stats package version 4.4.0. Only domains which were observed in 3 or more of our test set of Pangenes and with an enrichment P-value less than 0.05 were reported.

For the variation comparison between NLR pangenes and other pangenes, information about NLR-encoding genes and NLRs with highly variable coding regions in maize was obtained from Prigozhin et al. 2025 (21) and in soybean it was available via NLRCladeFinder (https://github.com/daniilprigozhin/NLRCladeFinder.git) that used the same method as described in Prigozhin et al. 2025 for gene annotation.

### Overview of K-PROB functions

K-PROB is a Python-based tool to detect candidate regulatory k-mer sequences associated with up- or down-regulation of gene expression from pan-genome and pan-transcriptome datasets. The workflow has two steps: 1) k-mer enumeration and clustering, and 2) mapping to identify k-mer clusters as putative CREs 3) k-mer-pangene filtering for functional regulatory sites (Fig. 2A). In the first step, K-PROB enumerates unique k-mer sequences from an input promoter FASTA file with a user-defined k value. Next, K-PROB orders k-mers by abundance and lexicographic rank to improve clustering robustness and reproducibility. Unique k-mers are then clustered with CD-HIT (68) using a similarity threshold 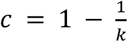, permitting one mismatch among unique k-mers within a cluster, and the following parameters: -r 1 -g 1 -gap -20 -gap-ext -10 -l -5 -M 0. Following k-mer clustering, K-PROB constructs a count matrix for “common” k-mer clusters, defined as clusters present in >1% of promoter sequences. This prevalence filter reduces matrix sparsity and model dimensionality, thereby increasing mapping power by lowering the number of parameters to estimate and stabilizing effect-size inference. After completing the count matrix construction, K-PROB leverages a Bayesian variable selection linear model, which is commonly used in polygenic score prediction (58, 69, 70), to map putatively causal k-mer sequences responsible for gene expression regulation.

Gene expressions are related to k-mer clusters with a standard linear regression model:

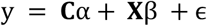

Where y is an n-dimensional vector of gene expression values, **C** is a N × c matrix of covariates, α is an n-dimensional vector of covariate effects, X is a N × p matrix of k-mer cluster count (**X**_**ij**_ denotes the count of k-mer cluster j in promoter sequence i), β is a p-dimensional vector of k-mer cluster effects, and ϵ is an n-dimensional vector of residual effects. Following a Bayesian hierarchical framework, K-PROB assumes:

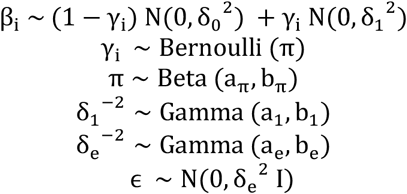

As shown in the model, K-PROB assumes the k-mer cluster effects follow a mixture of a normal density with mean 0 and variance δ_1_^2^, and a normal density with mean 0 and variance δ_0_^2^ with δ_0_^2^ ≪ δ_0_^2^. In practice, K-PROB sets δ_0_^2^ as 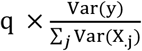, where q (default 0.05) is the proportion of the gene expression variation explained by the background effects (β |γ = 0). The latent variable γ encodes the component whose corresponding effects come from N(0, δ_0_^2^) or N(0, δ_1_^2^). An in-house Gibbs sampler was developed for parameter inference with the Geweke test for MCMC chain convergence (71). In the end, K-PROB outputs k-mer clusters with corresponding posterior inclusion probability (PIP) PIP_i_ = E(γ_i_| y, **C, X**), effect size, and false discovery rate estimates as metrics for prioritizing candidate causal k-mer sequences.

To further filter for functional sites, the effect of each k-mer cluster was assessed on a per-pangene basis by performing a simple linear regression between its occurrence and gene expression within each pangene. K-mer-pangene pairs were retained as putative functional sites only if they exhibited a significantly non-zero effect (FDR-corrected p < 0.05) and the estimated effect (i.e., the regression slope) was in the same direction as that predicted by the Bayesian model. This functional site filtering step was implemented using a custom R script.

### Using 10-fold cross validation to choose the best k value for K-PROB

Choosing k is critical for k-mer-based analyses. We treated k as a tunable hyperparameter and used 10-fold cross-validation to select the best value for the datasets. We partitioned the data into ten disjoint folds, trained K-PROB on nine folds to estimate k-mer cluster effects, and predicted gene expression in the held-out fold using those estimates. Each fold served once as the test set, ensuring all data were used for training and evaluation. We repeated this procedure for k from 7 to 12 and selected the k that yielded the highest mean prediction performance, evaluated by R^2^. To assess robustness, we tested three 10-fold partitioning schemes: (1) gene-level random splits, (2) pangene-level random splits, and (3) individual (accession)-level random splits. These schemes hide different sources of information during training and provide a comprehensive evaluation of k.

### TF binding motif alignment and enrichment analysis

Putative maize and soybean transcription factor binding motifs (TFBMs) were obtained from PlantTFDB (https://planttfdb.gao-lab.org/download.php#bind_motif). We tested whether the significant k-mer clusters identified by K-PROB were enriched for putative TF binding sequences using a permutation enrichment test. Specifically, for each permutation, we randomly sampled a set of k-mer clusters from the full set of clusters used in K-PROB. K-mers from these randomly sampled clusters were scanned against the TFBM database using FIMO v5.5.7 with the default significance threshold of p < 0.0001. We then counted how many clusters contained at least one k-mer with a significant match to a known TFBM. This process was repeated 10,000 times to generate a null distribution of the expected number of TFBM-matching clusters. The empirical p-value was calculated as the proportion of permutations in which the number of hit clusters exceeded that observed among the K-PROB significant clusters. All permutations and statistical analyses were performed in R using custom scripts. To evaluate whether motifs formed by significant k-mer clusters matched any known TFBMs, we first computed cluster motifs from the corresponding k-mer sequences weighted by their frequencies in promoters using MEME v5.5.7. The resulting motifs were then aligned to known TFBMs either from PlantTFDB or DAP-seq using TOMTOM v5.5.7 with an E-value cutoff of < 0.1. All permutations and statistical tests were implemented in R using custom scripts.

### Promoter luciferase reporter assay

*N. benthamiana* plants were grown in a Conviron growth chamber under 25°C/23 °C, a 14 h/10 h day/night cycle and 80 µmol/m^2^/s. Maize genomic DNAs were extracted from seed embryos using the Qiagen DNeasy Plant Kit (Qiagen, USA). 2kb maize promoters were cloned from the genomic DNAs using the Phusion DNA polymerase (New England Biolabs, USA) using primers with addition of the 5’ flanking overhang and a BsaI restriction site for downstream GoldenGate reactions (Dataset S5). Internal BsaI sites were removed using PCR-based single-site mutagenesis. Promoters were cloned into the fungal auto-bioluminescence pathway vector (kindly given by the Dr. Steinbrenner’s lab) for luciferase reporter assay via GoldenGate cloning (72, 73). Mutant promoters, in which only the target k-mer sequence was removed, were constructed by stitching together the flanking promoter fragments. *Agrobacterium tumefaciens* GV3101:pMP90 strains carrying the respective constructs were grown at 28°C in Luria-Bertani (LB) medium containing with 50 mg/ml Rifampicin, 25 mg/ml Gentamicin, and 50 mg/ml Kanamycin. Cultures were resuspended in infiltration medium (10 mM MES pH=5.6, 10 mM MgCl2 and 150 uM Acetosyringeone) at OD_600_=0.3. The top two fully expanded leaves of 4-5 weeks old N. benthamiana plants were infiltrated using a blunt syringe, with WT and mutant infiltrated on the same leaf to control the between-leaf expression efficiency variation. After two days post infiltration, luminescence was imaged with a ChemiDoc MP imaging system (Biorad, USA) using the “Chemiluminescence” function with optimal exposure times for each promoter. For quantification, the luminescence was measured using an Infinite F Plex Plate Reader (Tecan). Three leaf discs were measured from the same infiltration area as technical replicates and four biological replicates were used.

### Comparison to functional annotation datasets

Maize ATAC-seq narrow peak annotation file was obtained from NCBI GEO accession GSE165787 for all 26 lines (29), and Maize DAP-seq narrow peak annotation files of 212 TFs of B73 line were retrieved from Zenodo (https://doi.org/10.5281/zenodo.14991916) (16). An in-house python script was used to count the k-mer cluster occurrences in peaks and in all hv promoters. For each k-mer cluster, the percentage of occurrences overlapping with ATAC-seq peaks was calculated as the number of occurrences within narrow peaks divided by the total number of occurrences across all promoters. Similarly, the normalized k-mer occurrence at DAP-seq peaks was computed as the aggregated number of occurrences falling within DAP-seq narrow peaks from all available TFs, divided by the total number of occurrences across all promoters. All statistical tests and figure generation were implemented in R using custom scripts.

## Acknowledgments

We sincerely thank Dr. Daniil M. Prigozhin (Lawrence Berkeley National Laboratory) for providing the maize and soybean NLR gene annotations, Dr. George Chuck (UC Berkeley) for sharing maize inbred line materials, Dr. Miles Roberts (UC Berkeley) for valuable comments on the manuscript, and Dr. Li Lei (Syngenta) for constructive suggestions on the method. We also acknowledge the UC Berkeley Savio High-Performance Computing Cluster, its staff, and services for supporting our computational analyses. This work was supported by the National Institute of Health Director’s Award (1DP2AT011967-01) to Ksenia Krasileva.

## Data, Materials, and Software Availability

K-PROB is available via Github at https://github.com/xingwu2/K-PROB and other code was deposited at https://github.com/WeiWei-1112/K-RPOB-analysis-code-and-scripts.git. Processed data is available via Zenodo at doi.org/10.5281/zenodo.17346377. Previously published data containing the raw data used for this work was mentioned in materials and methods. All other data are included in the manuscript and/or supporting information.

## References

1. A. Hendelman, et al., Conserved pleiotropy of an ancient plant homeobox gene uncovered by cis-regulatory dissection. Cell 184, 1724–1739.e16 (2021).

2. J. A. N. Brophy, et al., Synthetic genetic circuits as a means of reprogramming plant roots. Science 377, 747–751 (2022).

3. R. J. Schmitz, E. Grotewold, M. Stam, Cis-regulatory sequences in plants: Their importance, discovery, and future challenges. Plant Cell 34, 718–741 (2022).

4. H. S. Mason, D. B. DeWald, J. E. Mullet, Identification of a methyl jasmonate-responsive domain in the soybean vspB promoter. Plant Cell 5, 241–251 (1993).

5. W. Wei, et al., Engineering pathogen-inducible promoters for conferring disease resistance in tomato. bioRxiv (2024).

6. X. Tu, et al., Reconstructing the maize leaf regulatory network using ChIP-seq data of 104 transcription factors. Nat. Commun. 11, 5089 (2020).

7. R. C. O’Malley, et al., Cistrome and epicistrome features shape the regulatory DNA landscape. Cell 165, 1280–1292 (2016).

8. K. A. Maher, et al., Profiling of Accessible Chromatin Regions across Multiple Plant Species and Cell Types Reveals Common Gene Regulatory Principles and New Control Modules. Plant Cell 30, 15–36 (2018).

9. A. P. Marand, Z. Chen, A. Gallavotti, R. J. Schmitz, A cis-regulatory atlas in maize at single-cell resolution. Cell 184, 3041–3055.e21 (2021).

10. B. Song, et al., Conserved noncoding sequences provide insights into regulatory sequence and loss of gene expression in maize. Genome Res. 31, 1245–1257 (2021).

11. S. Zenker, et al., Many transcription factor families have evolutionarily conserved binding motifs in plants. Plant Physiol. 198, kiaf205 (2025).

12. S. Inukai, K. H. Kock, M. L. Bulyk, Transcription factor-DNA binding: beyond binding site motifs. Curr. Opin. Genet. Dev. 43, 110–119 (2017).

13. T. Li, et al., Modeling 0.6 million genes for the rational design of functional cis-regulatory variants and de novo design of cis-regulatory sequences. Proc. Natl. Acad. Sci. U. S. A. 121, e2319811121 (2024).

14. F. F. Peleke, S. M. Zumkeller, M. Gültas, A. Schmitt, J. Szymański, Deep learning the cis-regulatory code for gene expression in selected model plants. Nat. Commun. 15, 3488 (2024).

15. J. Engelhorn, et al., Genetic variation at transcription factor binding sites largely explains phenotypic heritability in maize. bioRxiv 2023.08.08.551183 (2023).

16. M. Galli, et al., Transcription factor binding divergence drives transcriptional and phenotypic variation in maize. Nat. Plants 11, 1205–1219 (2025).

17. J. G. Wallace, et al., Association mapping across numerous traits reveals patterns of functional variation in maize. PLoS Genet. 10, e1004845 (2014).

18. W. Guo, et al., A barley pan-transcriptome reveals layers of genotype-dependent transcriptional complexity. Nat. Genet. 57, 441–450 (2025).

19. R. W. Michelmore, B. C. Meyers, Clusters of resistance genes in plants evolve by divergent selection and a birth-and-death process. Genome Res. 8, 1113–1130 (1998).

20. C. A. Sutherland, D. M. Stevens, K. Seong, W. Wei, K. V. Krasileva, The resistance awakens: Diversity at the DNA, RNA, and protein levels informs engineering of plant immune receptors from Arabidopsis to crops. Plant Cell 37 (2025).

21. D. M. Prigozhin, C. A. Sutherland, S. Rangavajjhala, K. V. Krasileva, Majority of the highly variable NLRs in maize share genomic location and contain additional target-binding domains. Mol. Plant. Microbe. Interact. 38, 275–284 (2025).

22. S. Ducrocq, et al., Key impact of Vgt1 on flowering time adaptation in maize: evidence from association mapping and ecogeographical information. Genetics 178, 2433–2437 (2008).

23. Y. Zan, X. Shen, S. K. G. Forsberg, Ö. Carlborg, Genetic regulation of transcriptional variation in natural Arabidopsis thaliana accessions. G3 (Bethesda) 6, 2319–2328 (2016).

24. K. A. G. Kremling, et al., Dysregulation of expression correlates with rare-allele burden and fitness loss in maize. Nature 555, 520–523 (2018).

25. X. Yuan, et al., Integrative omics analysis elucidates the genetic basis underlying seed weight and oil content in soybean. Plant Cell 36, 2160–2175 (2024).

26. N. Nariai, W. W. Greenwald, C. DeBoever, H. Li, K. A. Frazer, Efficient prioritization of multiple causal eQTL variants via sparse polygenic modeling. Genetics 207, 1301–1312 (2017).

27. Z. Zhang, et al., Major impacts of widespread structural variation on sorghum. Genome Res. 34, 286–299 (2024).

28. W. He, X. Li, Q. Qian, L. Shang, The developments and prospects of plant super-pangenomes: Demands, approaches, and applications. Plant Commun. 6, 101230 (2025).

29. M. B. Hufford, et al., De novo assembly, annotation, and comparative analysis of 26 diverse maize genomes. Science 373, 655–662 (2021).

30. Y. Liu, et al., Pan-genome of wild and cultivated soybeans. Cell 182, 162–176.e13 (2020).

31. L. Ni, et al., Pan-3D genome analysis reveals structural and functional differentiation of soybean genomes. Genome Biol. 24, 12 (2023).

32. E. K. Cannon, et al., Enhanced pan-genomic resources at the maize genetics and genomics database. Genetics 227, iyae036 (2024).

33. A. A. Kolodziejczyk, et al., Single cell RNA-sequencing of pluripotent states unlocks modular transcriptional variation. Cell Stem Cell 17, 471–485 (2015).

34. A. J. Aylward, S. Petrus, A. Mamerto, N. T. Hartwick, T. P. Michael, PanKmer: k-mer-based and reference-free pangenome analysis. Bioinformatics 39 (2023).

35. J. M. Ugalde, et al., A dual role for glutathione transferase U7 in plant growth and protection from methyl viologen-induced oxidative stress. Plant Physiol. 187, 2451–2468 (2021).

36. Y.-Q. Gao, et al., A dirigent protein complex directs lignin polymerization and assembly of the root diffusion barrier. Science 382, 464–471 (2023).

37. C. Paniagua, et al., Dirigent proteins in plants: modulating cell wall metabolism during abiotic and biotic stress exposure. J. Exp. Bot. 68, 3287–3301 (2017).

38. J. Yu, et al., A unified mixed-model method for association mapping that accounts for multiple levels of relatedness. Nat. Genet. 38, 203–208 (2006).

39. J. Baute, et al., Correlation analysis of the transcriptome of growing leaves with mature leaf parameters in a maize RIL population. Genome Biol. 16, 168 (2015).

40. D. Wilson-Sánchez, et al., Leaf phenomics: a systematic reverse genetic screen for Arabidopsis leaf mutants. Plant J. 79, 878–891 (2014).

41. X. Chen, Y. Hou, Y. Cao, B. Wei, L. Gu, A comprehensive identification and expression analysis of the WUSCHEL homeobox-containing protein family reveals their special role in development and abiotic stress response in Zea mays L. Int. J. Mol. Sci. 25, 441 (2023).

42. Z. Luo, et al., A dynamic regulome of shoot-apical-meristem-related homeobox transcription factors modulates plant architecture in maize. Genome Biol. 25, 245 (2024).

43. I.-C. Vélez-Bermúdez, et al., A MYB/ZML complex regulates wound-induced lignin genes in maize. Plant Cell 27, 3245–3259 (2015).

44. J. Jin, H. Zhang, L. Kong, G. Gao, J. Luo, PlantTFDB 3.0: a portal for the functional and evolutionary study of plant transcription factors. Nucleic Acids Res. 42, D1182–7 (2014).

45. K. de Silva, B. Laska, C. Brown, H. W. Sederoff, M. Khodakovskaya, Arabidopsis thaliana calcium-dependent lipid-binding protein (AtCLB): a novel repressor of abiotic stress response. J. Exp. Bot. 62, 2679–2689 (2011).

46. H. Zhang, Y. Zhao, J.-K. Zhu, Thriving under stress: How plants balance growth and the stress response. Dev. Cell 55, 529–543 (2020).

47. I. J. Higgins, S. G. Choudury, A. Y. Husbands, Mechanisms driving functional divergence of transcription factor paralogs. New Phytol. 247, 2022–2033 (2025).

48. A. Moons, Regulatory and functional interactions of plant growth regulators and plant glutathione S-transferases (GSTs). Vitam. Horm. 72, 155–202 (2005).

49. C. Vieira Dos Santos, P. Rey, Plant thioredoxins are key actors in the oxidative stress response. Trends Plant Sci. 11, 329–334 (2006).

50. L. Xu, et al., AIM1-dependent high basal salicylic acid accumulation modulates stomatal aperture in rice. New Phytol. 238, 1420–1430 (2023).

51. B. Mokhele, X. Zhan, G. Yang, X. Zhang, Review: Nitrogen assimilation in crop plants and its affecting factors. Can. J. Plant Sci. 92, 399–405 (2012).

52. M. D. Roberts, O. Davis, E. B. Josephs, R. J. Williamson, K-mer-based approaches to bridging pangenomics and population genetics. Mol. Biol. Evol. 42 (2025).

53. Y. Voichek, D. Weigel, Identifying genetic variants underlying phenotypic variation in plants without complete genomes. Nat. Genet. 52, 534–540 (2020).

54. Z. Zhang, et al., A k-mer-based pangenome approach for cataloging seed-storage-protein genes in wheat to facilitate genotype-to-phenotype prediction and improvement of end-use quality. Mol. Plant 17, 1038–1053 (2024).

55. D. Lee, R. Karchin, M. A. Beer, Discriminative prediction of mammalian enhancers from DNA sequence. Genome Res. 21, 2167–2180 (2011).

56. M. Ghandi, D. Lee, M. Mohammad-Noori, M. A. Beer, Enhanced regulatory sequence prediction using gapped k-mer features. PLoS Comput. Biol. 10, e1003711 (2014).

57. B. Schwarz, C. B. Azodi, S.-H. Shiu, P. Bauer, Putative cis-regulatory elements predict iron deficiency responses in Arabidopsis roots. Plant Physiol. 182, 1420–1439 (2020).

58. X. Wu, et al., Prioritized candidate causal haplotype blocks in plant genome-wide association studies. PLoS Genet. 18, e1010437 (2022).

59. W. R. Swindell, The association among gene expression responses to nine abiotic stress treatments in Arabidopsis thaliana. Genetics 174, 1811–1824 (2006).

60. S. Liu, et al., Mapping regulatory variants controlling gene expression in drought response and tolerance in maize. Genome Biol. 21, 163 (2020).

61. M. Kazemian, H. Pham, S. A. Wolfe, M. H. Brodsky, S. Sinha, Widespread evidence of cooperative DNA binding by transcription factors in Drosophila development. Nucleic Acids Res. 41, 8237–8252 (2013).

62. A. M. Bolger, M. Lohse, B. Usadel, Trimmomatic: a flexible trimmer for Illumina sequence data. Bioinformatics 30, 2114–2120 (2014).

63. A. Dobin, et al., STAR: ultrafast universal RNA-seq aligner. Bioinformatics 29, 15–21 (2013).

64. B. Li, C. N. Dewey, RSEM: accurate transcript quantification from RNA-Seq data with or without a reference genome. BMC Bioinformatics 12, 323 (2011).

65. A. R. Quinlan, I. M. Hall, BEDTools: a flexible suite of utilities for comparing genomic features. Bioinformatics 26, 841–842 (2010).

66. S. B. Cannon, H.-O. Lee, N. T. Weeks, J. Berendzen, Pandagma: a tool for identifying pangene sets and gene families at desired evolutionary depths and accommodating whole-genome duplications. Bioinformatics 40 (2024).

67. H. Einarsson, et al., Promoter sequence and architecture determine expression variability and confer robustness to genetic variants. Elife 11 (2022).

68. L. Fu, B. Niu, Z. Zhu, S. Wu, W. Li, CD-HIT: accelerated for clustering the next-generation sequencing data. Bioinformatics 28, 3150–3152 (2012).

69. T. Ge, C.-Y. Chen, Y. Ni, Y.-C. A. Feng, J. W. Smoller, Polygenic prediction via Bayesian regression and continuous shrinkage priors. Nat. Commun. 10, 1776 (2019).

70. X. Zhou, P. Carbonetto, M. Stephens, Polygenic modeling with bayesian sparse linear mixed models. PLoS Genet. 9, e1003264 (2013).

71. J. Geweke, Evaluating the accuracy of sampling-based approaches to the calculation of posterior moments. Staff Report (1991).

72. A. G. K. Garcia, A. D. Steinbrenner, Bringing Plant Immunity to Light: A Genetically Encoded, Bioluminescent Reporter of Pattern-Triggered Immunity in Nicotiana benthamiana. Mol. Plant. Microbe. Interact. 36, 139–149 (2023).

73. E. Weber, C. Engler, R. Gruetzner, S. Werner, S. Marillonnet, A modular cloning system for standardized assembly of multigene constructs. PLoS One 6, e16765 (2011).

